# Oxygen as a primary selective pressure for photo-endosymbiosis evolution

**DOI:** 10.1101/2025.05.28.656558

**Authors:** Loïc Quevarec, Rachel Bonnarde, Christophe Robaglia, Gaël Brasseur

## Abstract

Photosymbiosis through eukaryotic or eukaryotic – prokaryotic associations provide mainly carbon and oxygen to the living world, however the mechanisms behind their evolution remain unknown. Here we develop a naïve system based on the predatory ciliate *Tetrahymena thermophila*, not known for hosting symbiont, to recapitulate early events leading to photosymbiosis evolution. *T. thermophila* was found to readily phagocyte eukaryotic algae (*Chlorella variabilis*) or cyanobacteria (*Synechoccocus elongatus*) that can reach high numbers within predatory cells. Feeding with either prey in a low carbon environment provide little or no growth advantage to *T. thermophila*. In contrast, in anoxic environment, both intracellular *C. variabilis* or *S. elongatus* can readily support growth of *T. thermophila* in the light but not in the dark, demonstrating that oxygen supply *in hospite* can be an initial feature supporting further photo-endosymbiosis evolution. This suggests that it is unlikely that carbon metabolites produced by phototrophs could have represented a selective advantage for the evolution of photosymbionts. Although most extant photosymbiosis are based on carbon supply to the host cell, we therefore propose that it is a secondary event occurring from initial evolution in anoxic or hypoxic conditions, where O_2_ production was crucial for establishing the initial steps of photosymbiosis.

## Introduction

Oxygenic photosynthesis appeared in cyanobacteria probably around 3 Byr ago (Fischer et al., 2016). Using light energy, it provides the chemical energy allowing to fix atmospheric CO_2_ into building block of life, supporting the proliferation of photosynthetic organisms and indirectly the organisms that feed on them. Photosynthesis was acquired by eukaryotes by engulfing cyanobacteria, around 1.6 Byr ago (Archibald, 2015; Bhattacharya et al., 2023), leading to the Archaeplastida group containing the Glaucophyta, the Rhodophyta (red algae) and the Viridiplantae (green algae and land plants) (Ponce-Toledo et al., 2019). Integration of a photosynthetic prokaryote within a eukaryote is known as primary endosymbiosis and has also been described for the amoeba *Paulinella chromatophora* and dated around 90–140 million years ago (Delaye et al., 2016; Ponce-Toledo et al., 2019). Further spreading of photosynthesis occurred by integration of photosynthetic eukaryotes within eukaryotes and are known as secondary and higher order endosymbiosis events. They concern the majority of eukaryotic groups (Lane et Archibald, 2008; Ponce-Toledo et al., 2019). Photosynthetic endosymbiosis is globally present on the entire surface of the Earth and have a considerable impact on environments (O_2_ production, CO_2_ and N_2_ fixation, …) and the other species that populate them (Bowles et al., 2023).

The recurrent acquisition of photosynthesis by photosymbiosis across the tree of life (Ponce-Toledo et al., 2019), indicates that it is a highly advantageous trait, selected multiple times independently, i.e. convergent evolution (Keeling, 2010). Observable cases of photosymbiosis show that symbionts are initially acquired by predation, particularly by phagocytosis (Gavelis et Gile, 2018; Inouye et Okamoto, 2005; Quevarec et al., 2024). The production of sugars or oxygen by the photosynthetic symbiont and supplied to the host are likely to support the evolution of photosymbiosis (Germond et al., 2013). Extant stable photosymbiosis in plants and algae are clearly based on carbon autotrophy (Hoefnagel et al., 1998). In the other hand, for example, the photosymbiotic ciliate *Tetrahymena utriculariae* lives in the anoxic traps of the aquatic carnivorous plant *Utricularia reflexa*, where dissolved carbon and other nutrients are abundant (Pitsch et al., 2017). However, most data, if not all, come from already established photosymbiotic associations, where host and symbiont had ample time to co-evolve, therefore the evolutionary trajectories and early events leading to stabilised associations remain unknown.

In this work, using a predator-prey model system, we experimentally evaluate what could be the primary selective force for photosymbiosis establishment. We used the ciliate *T. thermophila* as a model of predatory unicellular host, the cyanobacterium *Synechococcus elongatus* and the eukaryotic green algae *Chlorella variabilis* as preys. Those can recapitulate early events leading to either primary (*S. elongatus*) or secondary (*C. variabilis*) endosymbiosis. *T. thermophila* is a widely used model organism, that can grow by osmotrophy or phagocytosis of various preys (Lynn et Doerder, 2012; Wloga et Frankel, 2012). Despite being studied since many years, it has never been found to stably host photosynthetic symbionts in the wild. However, related species such as *Paramecium bursaria* (Fujishima et Kodama, 2022) or *T. utricularia* (Pitsch et al., 2017) and many other ciliates are known to host algal symbionts (Johnson, 2011). The freshwater cyanobacterium *S. elongatus* is a well described photosynthetic species (Adomako et al., 2022), living as a possible symbiont within *Crassostrea gigas* tissues (Avila-Poveda et al., 2014). The *Synechococcus* genus has the greatest genetic proximity to the *P. chromatophora* chromatophore which results from a recent primary endosymbiosis (Delaye et al., 2016; Nowack et al., 2008; Ponce-Toledo et al., 2019). *C. variabilis* is a well described photosynthetic algal species (Krienitz et al., 2015), which together with others members of its genus, are symbiont in many endosymbiosis (i.e. with *P. bursaria* (Zagata et al., 2016), *Hydra viridissima* (Rajević et al., 2015), *Stentor pyriformis* (Hoshina et al., 2021) or *Acanthocystis turfacea* (Matzke et al., 1990)).

We show that in an initial predatory-prey state, the supply of carbon by photosynthetic prey support modestly the growth of the predator, while the supply of oxygen leads to a readily observable growth advantage, making it a likely proximal selective pressure for initiation of photo-endosymbiosis.

## Results

### Phagocytosis dynamics

In coculture of cyanobacterium *S. elongatus* UTEX 2973 (Fig. 1B) or the algae *C. variabilis* (Fig. 1C) with *T. thermophila* (Fig. 1A), this latter multiplied by asexual division and the resulting two daughter cells share the preys during mitosis showing a vertical transmission of the preys through generations (Fig. 1D and E). In presence of the cyanobacterium *S. elongatus*, *T. thermophila* phagosomes containing several red autofluorescent bacteria were readily observed (Fig. 1F). Flow cytometry using both scatter and chlorophyll-fluorescence signals was further used to quantify the rate and the amplitude of the phagocytosis process. This experiment showed a fast period of bacterial phagocytosis leading to a maximum of 160 cyanobacteria accumulated per *Tetrahymena* cells at a calculated maximum initial speed of about 800 cyanobacteria phagocyted per hour (1 cyanobacteria phagocyted every 4.5 s) (Fig. S2B). This period is followed by a longer period during which cyanobacterial cells progressively disappeared at a rate of about 20 cyanobacteria per hour (Fig. 2B). This kind of phagocytic behaviour was previously reported for *Tetrahymena pyriformis* (Thurman et al. 2010). We noticed that around 40 min after the initiation of coculture, a new population of fluorescent particles appears with an intermediate size between *T. thermophila* and *S. elongatus*. Fluorescence microscopy shows that they were often localized at the cytoproct and were droppings (faecal pellets) ejected by *Tetrahymena* and containing up to 4 cyanobacterial cells (Fig. 2A, C, D and Fig. 1G for droppings image).

**Figure 1:**
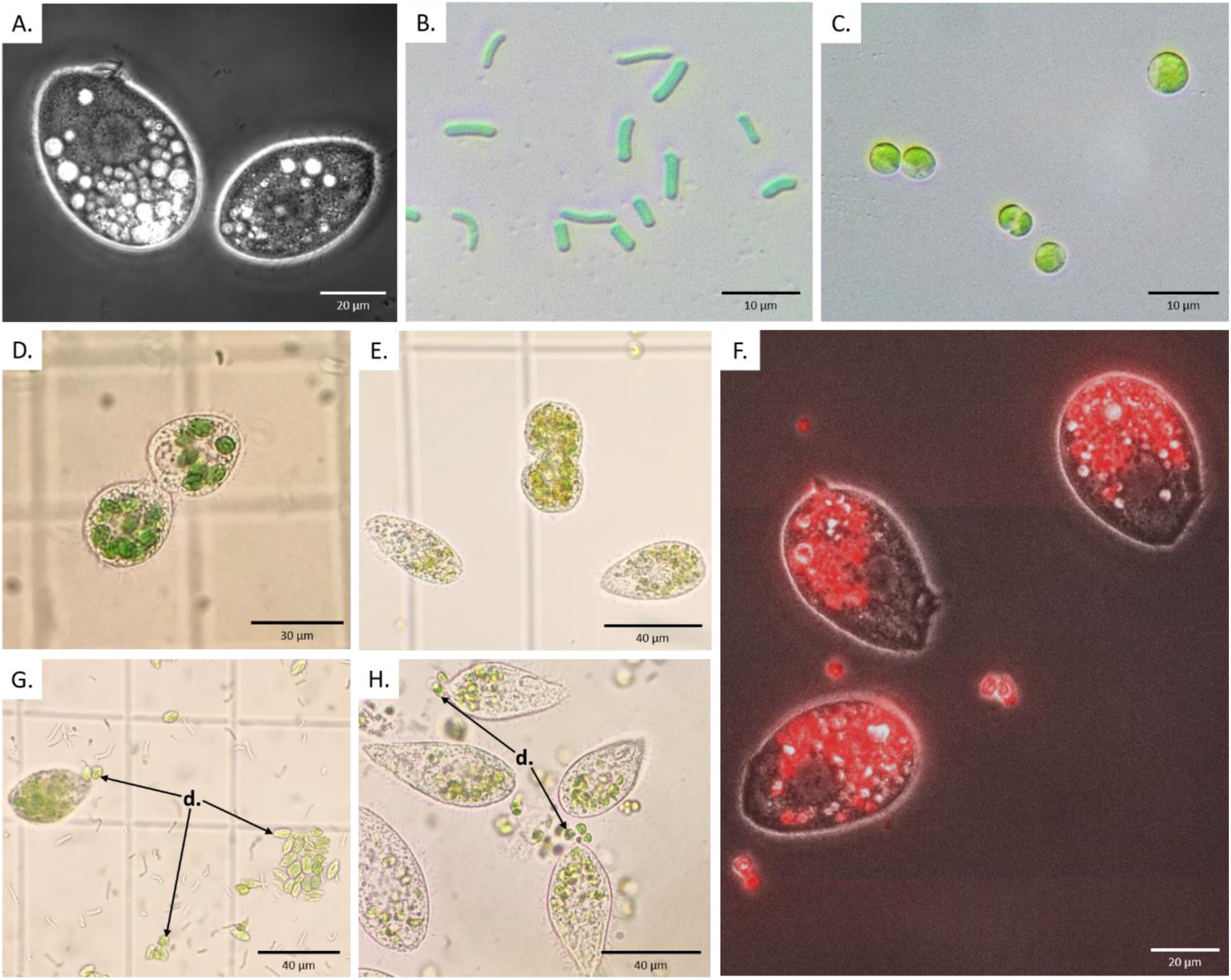
Optical microscopy representing A. *T. thermophila*, B. *S. elongatus* and C. *C. variabilis* cells. **D.** and **E.** *S. elongatus* and *C. variabilis* in *T. thermophila* cells undergoing mitosis showing vertical transmission, respectively. **F.** *C. variabilis* phagocytosed by *T. thermophila* shown by red fluorescence. **G.** and **H.** Green droppings (**d.**) exocytosed by *T. thermophila* and containing *S. elongatus* and *C. variabilis* cells, respectively.

**Figure 2:**
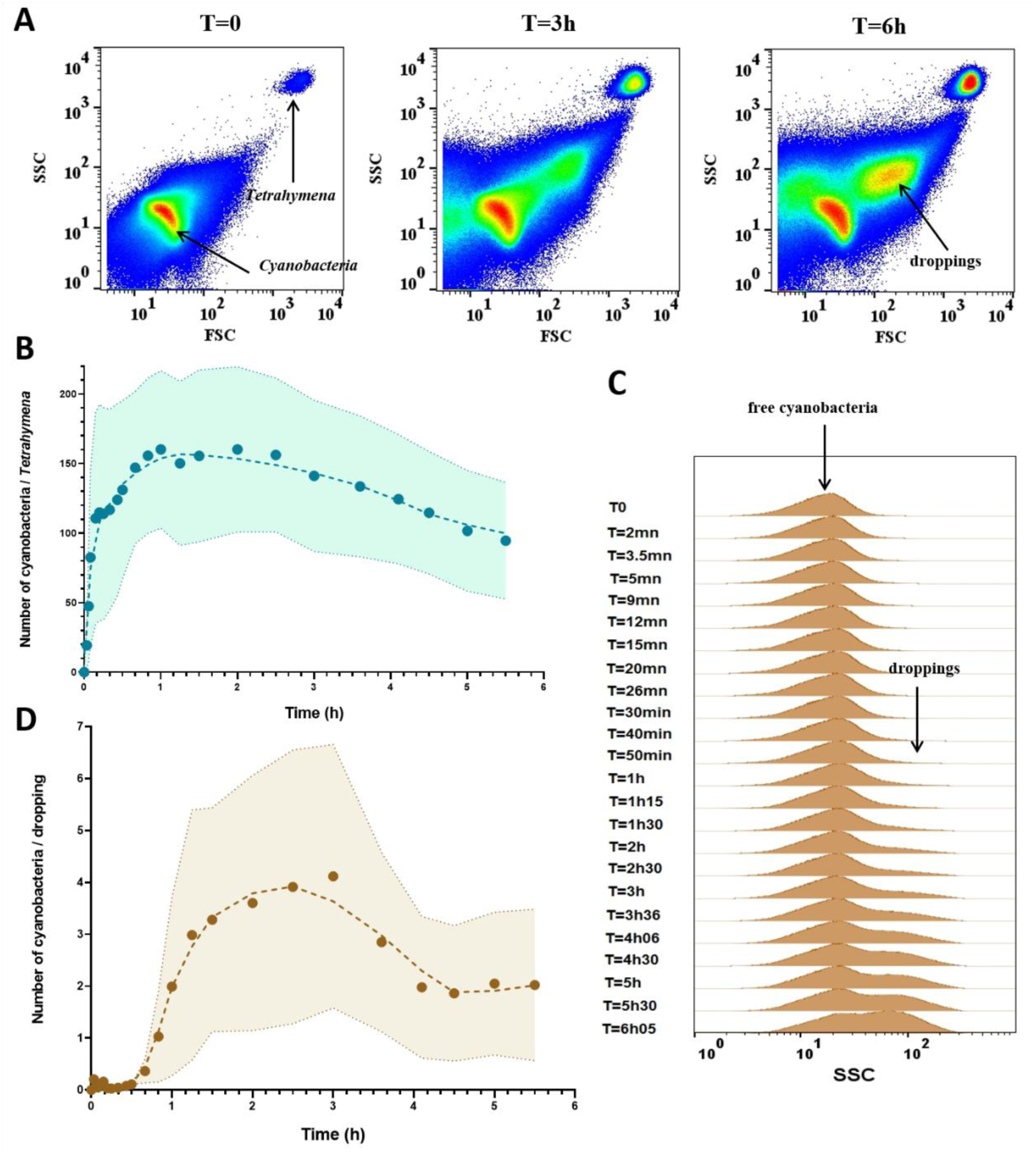
Phagocytosis of cyanobacteria *S*. *elongatus* (SE) by the ciliate *T*. *thermophila* (TT) **A)** Analysis of populations of cyanobacteria SE mixed with TT in oxic conditions at 30°C (initial ratio of 400 SE/TT). The apparition of TT droppings composed of fluorescent SE is indicated. Color code from blue to red indicates low to high relative density of populations respectively, in a constant number of total events per time point (n = 8 x10^5^). **B)** Kinetics SE transit inside of TT using the intrinsic red autofluorescence of SE. Shaded area corresponds to the standard deviation of fluorescence mean (n>10^3^). **C)** SSC scatter histograms of free cyanobacteria SE and the new population corresponding to the formation of droppings (see Fig. 2A). **D)** Kinetics of droppings formation expressed as the number of SE per dropping as a function of time.

In a contact between *T. thermophila* and *C. variabilis*, a first rapid phase of phagocytosis at a maximum rate of about 40 algae phagocyted per hour (1 every 1.6 min) succeeded a slower phase where *C. variabilis* cells progressively disappeared at a rate of 3.7 cells per hour (Fig. 3A, B and Fig. S3; Movie S1). The average size of free remaining *C. variabilis* cells increased during the phagocytosis process, suggesting a selective preference by *T. thermophila*for smaller *C. variabilis* possibly linked to the size of its oral apparatus (6-10 µM) compared to that of *C. variabilis* cells (3-10 µM) (Fig. S3B).

**Figure 3:**
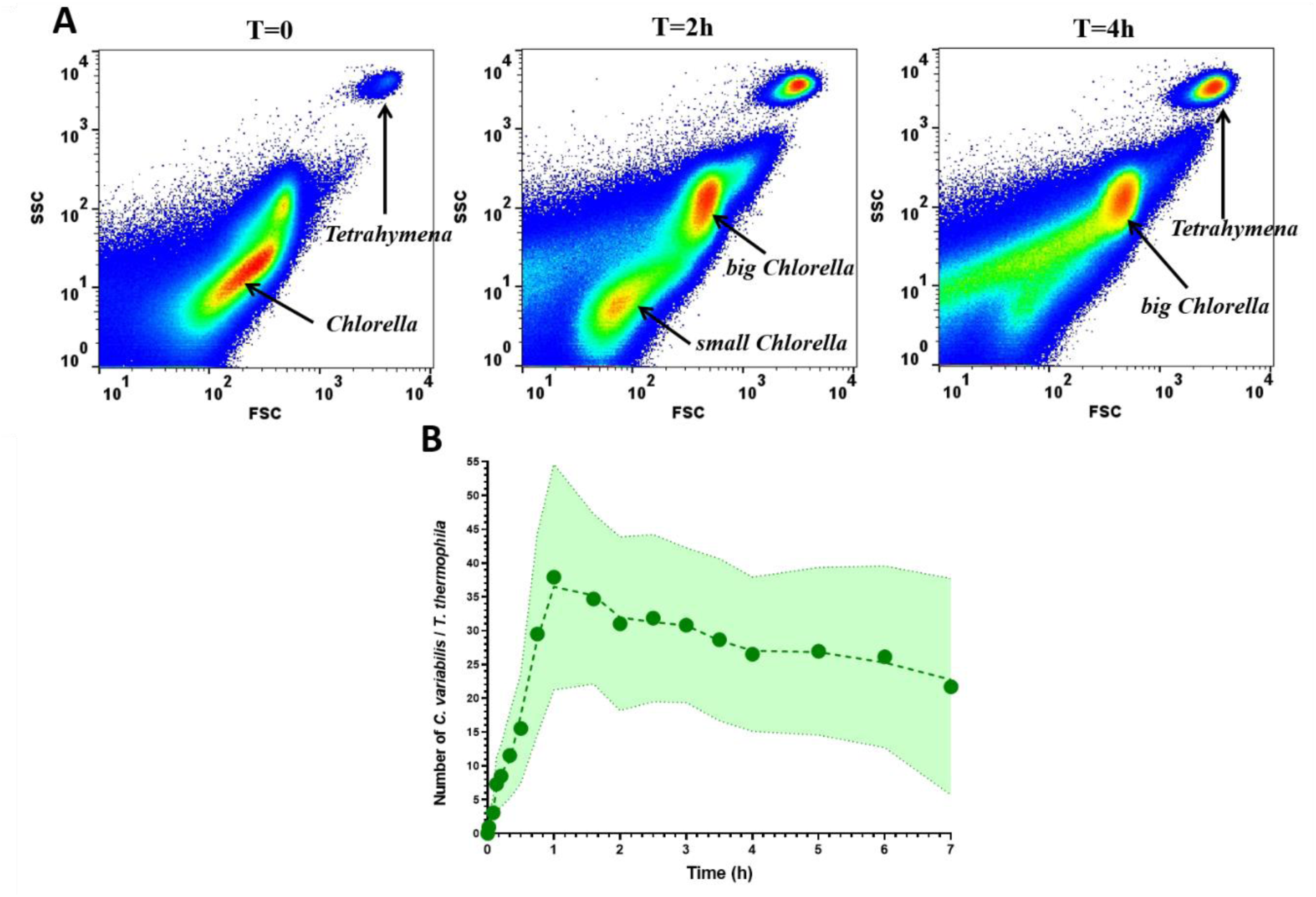
Phagocytosis of *C. variabilis* (CV) by *T. thermophila* (TT). **A)** Analysis of populations of Chlorella mixed with TT in oxic conditions at 30°C (initial ratio of 200 CV/TT). Flow cytometry scatter density plots (SSC versus FSC) shows the larger size of TT, the smaller diffraction of the CV and the disappearing of smaller CV as a function of time. Color code from blue to red indicates low to high relative density of populations respectively, in a constant number of total events per time point (n = 8 x 10^5^). **B)** Kinetics of the transit time of CV phagocyted per TT using the intrinsic red autofluorescence of CV. Shaded area corresponds to the standard deviation of fluorescence mean (n>10^4^).

In the presence of a GFP labelled *Escherichia coli* bacterium, up to 700 bacterial cells were phagocyted in one hour with a maximum initial rate at the onset of more than 3000 *E. coli*/hour (Fig. S4B – D). After one hour of phagocytosis, droppings containing up to 7 *E. coli*/dropping appeared (Fig. S4A and E).

In the presence of both *E. coli*-GFP and *C. variabilis*, *T. thermophila* ingested both type of cells simultaneously. *E. coli* phagocytosis was found markedly faster than that of *C. variabilis,* possibly linked to its smaller size (data not shown and compare Fig. 3B with Fig. S4C).

### Carbon benefit for *T. thermophila* through phagocytosis of phototrophs

We tested the contribution of carbon (sugars) produced by photosynthetic preys on the growth of *T. thermophila. T. thermophila* grew significantly in the rich Neff medium with *S. elongatus* (Fig. 4A; Table S1c) or *C. variabilis* (Fig. 4B; Table S2c) or without phototrophs compared with a culture of *T. thermophila* in mineral minimum TAP medium (Fig. 4A and B; Table S1c and S2c). In this rich Neff medium, the presence of phototrophs did not increase the growth yield and rate of *T. thermophila* (Fig. 4A and B, Table S1c and S2c). In the mineral minimum TAP medium, no growth of *T. thermophila* alone was observed even with the addition of *C. variabilis* (Fig. 4B; Table S2c). However, adding *S. elongatus* increased slightly the growth of *T. thermophila* (Fig. 4A; Table S1c).

**Figure 4:**
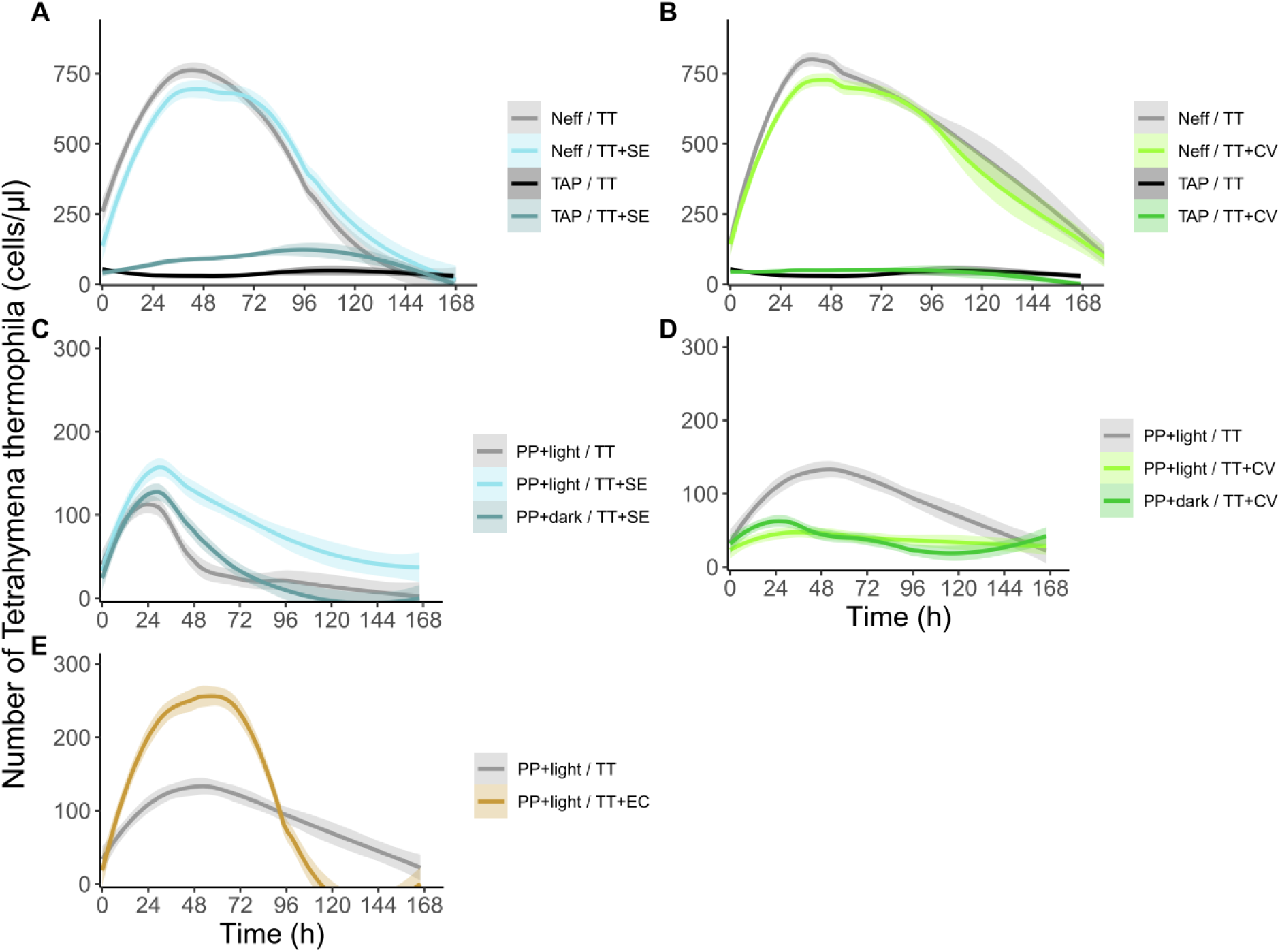
Contribution of prey microorganisms as carbon sources for *T. thermophila* (TT) growth. Shaded areas represent 95% confidence intervals. **A.** and **B.** Growth of *T. thermophila* in rich Neff medium and mineral minimum TAP medium with **A.** *S. elongatus* (SE) or **B.** *C. variabilis* (CV) as prey; **C.** and **D.** Growth in modified minimum PP medium in light or dark with **C.** *S. elongatus* (SE) or **D.** *C. variabilis* (CV); **E.** Growth in modified minimum PP medium with *E. coli* (EC). The data were analyzed using GAMM. All the results of the statistical test are shown in Table S1 - S5. Briefly, we observed a significant growth of *T. thermophila* in the rich Neff medium with SE (**A.** p-value < 0.001***) or CV (**B.** p-value < 10^-6^***) or without phototrophs (**A.** p-value < 0.001*** and **B.** p-value < 10^-6^***) compared with a culture on minimum TAP medium. In TAP medium, only SE increased slightly the growth of TT (**A.** p- value < 0.001***). In the modified minimum PP medium, **C.** SE stimulated the growth of TT in the light (p-value = 5.36 x 10^-5^***) but not in the dark (p-value = 0.556), while **D.** CV decreased the growth of TT both in the light (p-value = 0.000324***) and in the dark (p-value = 0.000376***). **E.** *E. coli* (EC) stimulated the growth of TT (p-value = 5.69 x 10^-8^***). ns: no significant; *, *P* < 0.05; **, *P* < 0.01; ***, *P* < 0.001.

In the modified PP minimum medium (Fig. 4C – E), the growth of *T. thermophila* is stimulated in the presence of *S. elongatus* in the light but not in the dark (Fig. 4C; Table S3a). With *C. variabilis*, a significant decrease in the growth of *T. thermophila* both in the light and in the dark was observed compared with a culture of *T. thermophila* alone (Fig. 4D; Table S4a). The non-photosynthetic *E. coli* significantly fuelled the growth of *T. thermophila* in PP minimum medium (Fig. 4E; Table S5a), although this growth is still less efficient than *T. thermophila* alone in the rich Neff medium (Fig. 4A).

### Oxygen benefit for *T. thermophila* through phagocytosis of phototrophs

We probed the lower limit of oxygen concentration to allow *T. thermophila* growth (Fig. 5A). Optimum growth in continuous oxygen bubbling occurs at atmospheric tension of 21% O_2_, while significant growth occurs with as little as 0.05% oxygen compared with a totally anoxic environment (Fig. 5A; Table S6).

**Figure 5:**
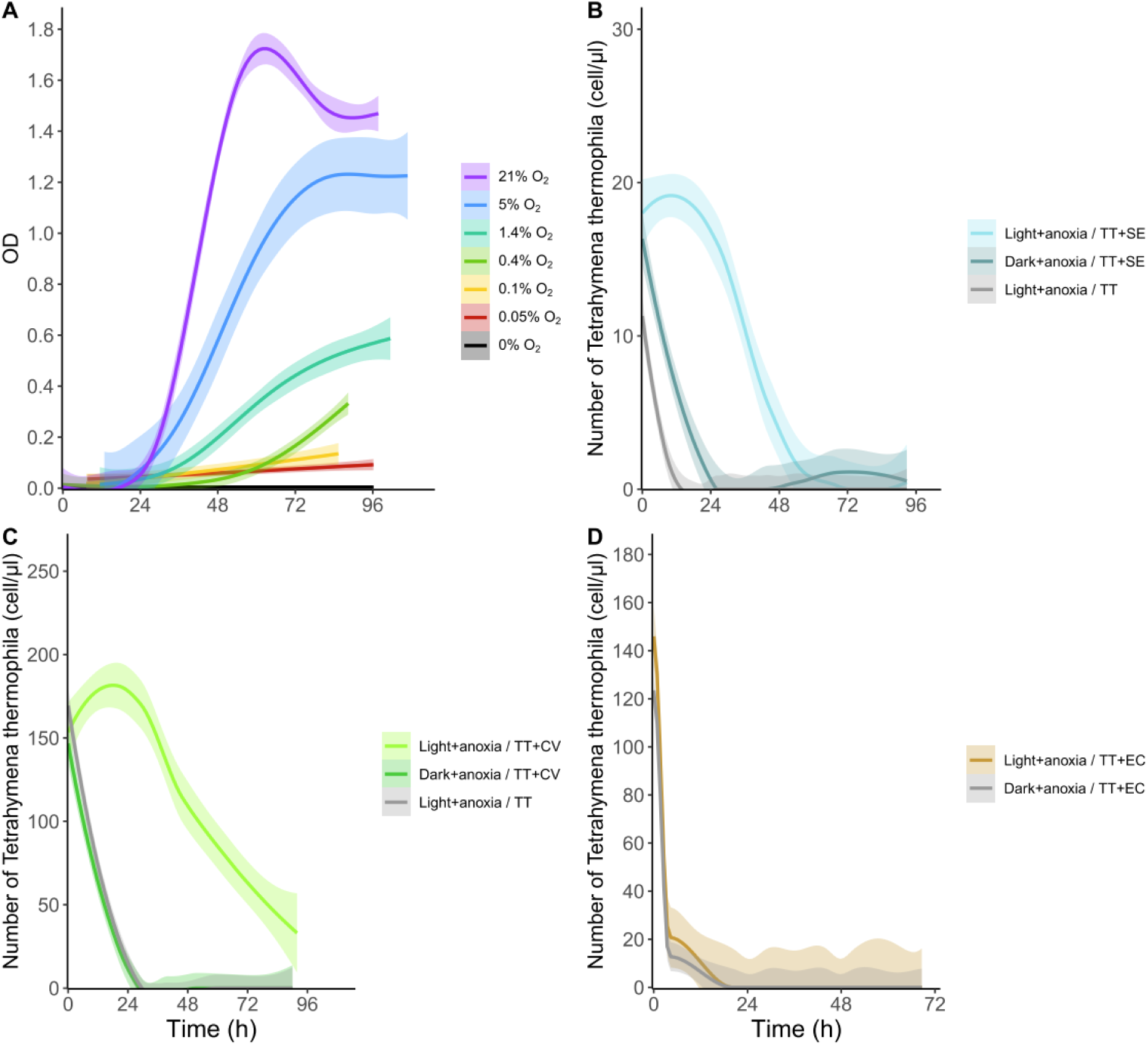
Contribution of prey microorganisms to *T. thermophila* (TT) growth in limiting oxygen. Shaded areas represent 95% confidence intervals. **A.** Growth of *T. thermophila* by measuring the OD (λ = 600 nm) of the culture with increasing oxygen concentrations ranging from 0% (pure N_2_) to atmospheric level of 21% O_2_. The data were analyzed using a GAMM. Shaded areas represent 95% confidence intervals. All the results of the statistical test are shown in Table S6. Briefly, we observed a significant growth for 0.05% oxygen (p-value = 0.0138*), 0.1% (p-value = 0.0119*), 0.4% (p-value = 0.011*), 1.4% (p-value = 5.22 x 10^-6^***), 5% (p-value = 5.27 x 10^-12^***), 21% (p-value < 2 x 10^-16^***). **B.** to **D.** Growth of *T. thermophila* under anoxic conditions in light or dark, with or without prey (**B.** *S. elongatus* (SE), **C.** *C. variabilis* (CV) and **D.** *E. coli* (EC)). The data were analyzed using a Kruskal-Wallis rank sum test and a post hoc pairwise comparison (Dunn’s test) with Holm correction. Results: **B.** Kruskal-Wallis chi-squared = 68.647, df = 2, p-value = 1.24 x 10^-15^***: light + TT vs light +TT + SE: Dunn’s test: adjusted p-value = 5.43 x 10^-16^***, dark +TT +SE vs light + TT + SE: Dunn’s test: adjusted p-value = 2.78 x 10^-5^***. **C.** Kruskal-Wallis chi-squared = 106.93, df = 2, p-value < 2.2 x 10^-16^***: light + TT vs light + TT + CV: Dunn’s test: adjusted p-value = 4.31 x 10^-13^***, dark + TT + CV vs light + TT + CV: Dunn’s test: adjusted p-value = 3.29 x 10^-20^***. **D.** dark +TT + EC vs light + TT + EC: Kruskal-Wallis chi-squared = 0.3375, df = 1, p- value = 0.5613. ns: no significant; *, *P* < 0.05; **, *P* < 0.01; ***, *P* < 0.001.

We then initiate anoxic cocultures in conditions where most phototrophs were found to be phagocyted within a short period of time, as defined both by flow cytometry and confirmed by fluorescent microscopy. In the dark and anaerobic conditions, *T. thermophila* did not grow and even lysed in the presence of all kinds of preys (Fig. 5B and C). In contrast phototrophs efficiently support *T. thermophila* growth in anoxia significantly in the light but not in the dark. In anoxic coculture with the non-phototrophic *E. coli, T. thermophila* did not grow either in the light or in dark (Fig. 5D).

These experiments strongly suggest that oxygen evolved by phagocyted photosynthetic preys in the light can rescue *T. thermophila* growth in anoxia. This was further supported in the experiments where *T. thermophila* and phototrophs cultures were separated by a tubing and sterile gas filter that only allow gaseous products to diffuse between flasks (data not shown).

We next evaluated the influence of anoxia on the phagocytosis competence of *T. thermophila* using flow cytometry. In anoxia in the dark, *T. thermophila* is unable to phagocyte either *S. elongatus* or *C. variabilis* whereas in anoxia and in the light, the phagocytosis is restored up to more than 100 *S. elongatus* and about 10 *C. variabilis* per *T. thermophila* cell as compared with oxic conditions (Fig. 6A, B). The phagocytosis and the transit time of ingested photosynthetic cells was found to be reduced in anoxia and light compared to oxic conditions (Fig. 6A and B).

**Figure 6:**
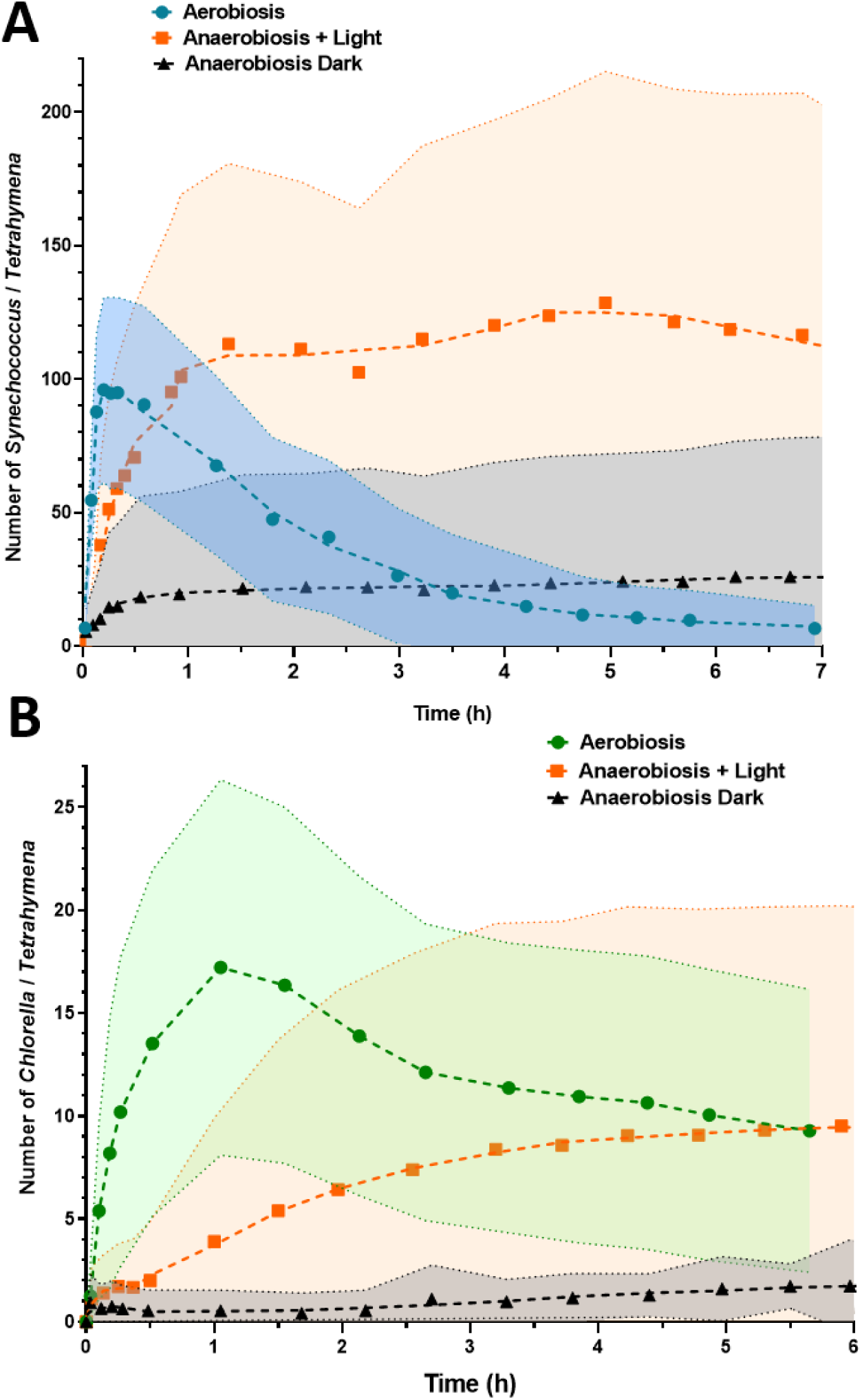
O_2_ produced by the phototrophs allows *T. thermophila* (TT) phagocytosis in anaerobiosis. **A)** Kinetics of *S. elongatus* (SE) transit within TT in oxic condition (atmospheric 21% O_2_, cyan dots), in anoxic conditions in the dark (black triangles) or in anoxic conditions in the light (orange squares) at 30°C (initial ratio of 500 SE/TT). **B)** Kinetics of *C. variabilis* (CV) transit time within TT in oxic condition (atmospheric 21% O_2_, green dots), in anoxic conditions in the dark (black triangles) or in anoxic conditions in the light (orange squares) (initial ratio of 150 CV/TT). Shaded area corresponds to the standard deviation of fluorescence mean (total events per time point: n = 10^6^).

### Prey O_2_ production and predator O_2_ consumption are compatible

We measured O_2_ consumption by *S. elongatus, C. variabilis* and *T. thermophila* and O_2_ production by *S. elongatus* and *C. variabilis* at their maximal rate in presence of saturating amount of substrate (glucose) or light with a modified oxymeter device (Fig. 7; Table 1; Fig. S5). *T. thermophila* mitochondrial O_2_ consumption amounts to 43 nmol O_2_/min/mg protein (i.e. 430 x 10^-7^ nmoles O_2_/min/cell, calculated with the calibration curves in Fig S1) with endogenous substrates present in the cells (Fig. S5B) and to 60 nmol O_2_/min/mg (i.e. 600 x 10^-7^ nmoles O_2_/min/cell) using glucose as substrate (Fig. 7; Table 1). This activity is completely inhibited by cyanide, a terminal oxygen reductase inhibitor (Fig. S5B – C). *C. variabilis* and *S. elongatus* exhibits an O_2_ consumption activity in the dark of 67 and 20 nmol O_2_/min/mg, respectively (Fig. 7 and S5C – D). For O_2_ production in the light, *C. variabilis* and *S. elongatus* produced 190 and 105 nmol O_2_/min/mg (i.e. around 2.3 and 0.4 x 10^-7^ nmoles O_2_/min/cell) (Table 1; Fig. S5C – D), respectively. This implies that about 260 *C. variabilis*. and 1500 *S. elongatus* per *T. thermophila* cell would be required to provide maximum O_2_ consumption by *T. thermophila* at atmospheric oxygen concentration (21% O_2_), while only a fraction of these numbers would be required to provide the minimal O_2_ concentration supporting *T. thermophila* growth (0.05% O_2_). After 60 min of phagocytosis, *S. elongatus* in phagosomes are still able to produce O_2_ with light at a similar rate compared to free cyanobacteria (Fig. S5D). Oxygen production by the phototrophs preys inside the host is quantitatively compatible with oxygen consumption by the ciliate host organism.

**Figure 7:**
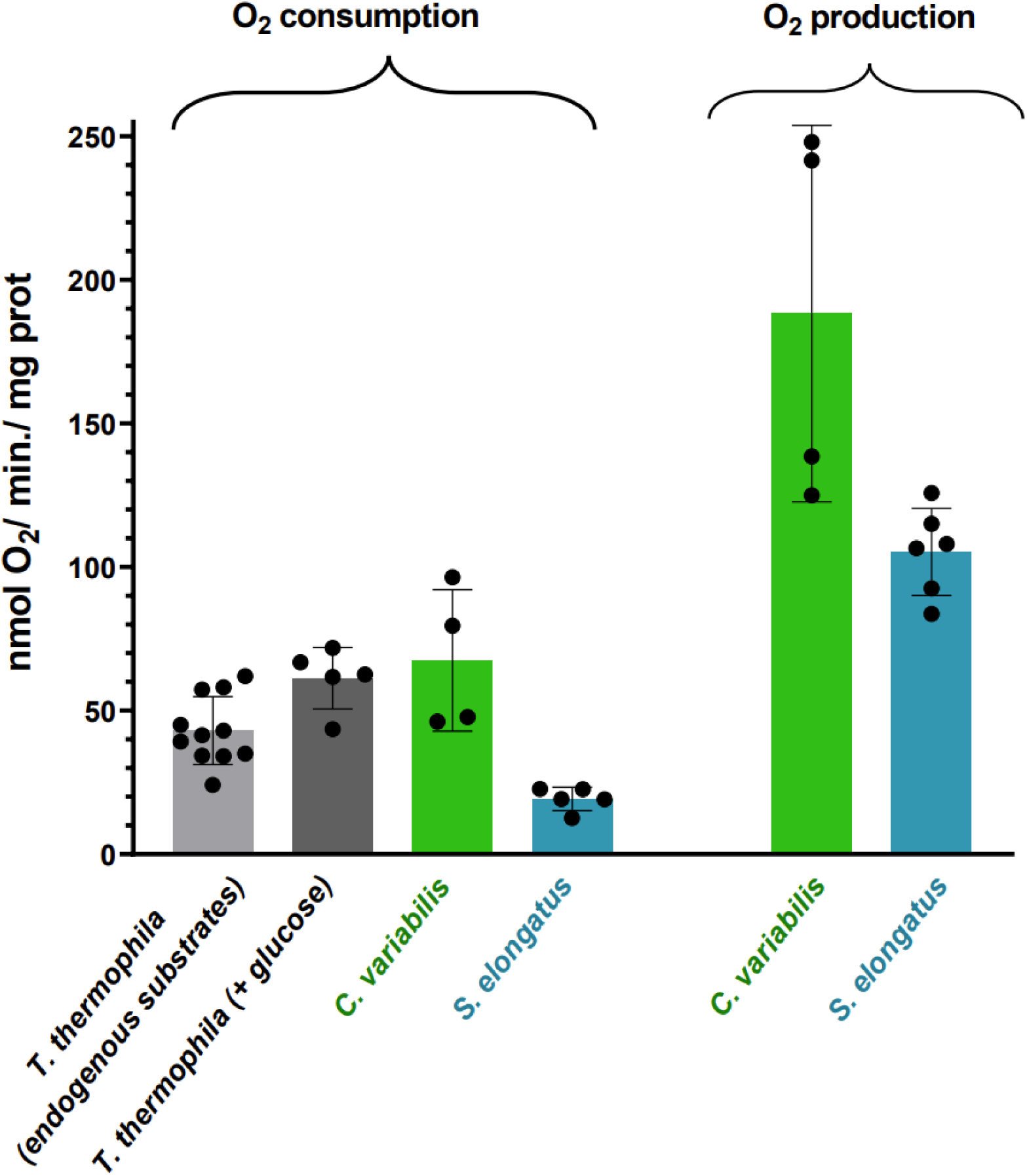
O_2_ evolution by the photosynthetic preys cyanobacteria and chlorella is compatible with *T. thermophila* aerobic metabolism. O_2_ consumption and production activities (nmol O_2_/mg/min, see Fig. S5) corresponding to maximum values with endogenous substrates, in saturation of substrate (glucose 20 mM) or in presence of saturating light for the phototrophs (60-100 µmoles photons/m^2^/s) are shown for the three organisms. See Table 1 for production and consumption of O_2_ at the single cell level.

**Table 1:** O_2_ evolution by the phototrophs cyanobacteria and chlorella and O_2_ consumption by *T. thermophila* at the single cell level are compatible with photo-endosymbiosis. From calibration curves presented in Fig. S6, O_2_ production and consumption at the cellular level for the three organisms were calculated. Results from Fig. 7 expressed per cell indicate that for maximum *T. thermophila* oxygen consumption rate, it requires the equivalent of about 260 chlorella and 1500 cyanobacteria producing O_2_ by photosynthesis.

| | $\text{O}_2$ consumption<br>(nmol $\text{O}_2$/min./cell) | $\text{O}_2$ production<br>(nmol $\text{O}_2$/min./cell) | *T. thermophila* $\text{O}_2$ consumption /<br>phototroph $\text{O}_2$ production |
| --- | --- | --- | --- |
| *Tetrahymena thermophila* | $600 \times 10^{-7}$ | n.a. | n.a. |
| *Chlorella variabilis* | $0.82 \times 10^{-7}$ | $2.3 \times 10^{-7}$ | 260 |
| *Synechococcus elongatus* | $0.073 \times 10^{-7}$ | $0.40 \times 10^{-7}$ | 1500 |

## Discussion

Stable photosymbiosis evolved independently in many species of ciliates, including *Tetrahymena utriculariae*. *T. thermophila* is not known to host photosynthetic symbionts in nature but is able to phagocyte different species of prokaryotic or eukaryotic phototrophs and long-term co-culture with algae and bacteria has been described (Germond et al., 2013; Sano et al., 2009). Assuming that photosymbiosis evolved from predation, this makes *T. thermophila* a suitable evolutionary model to explore the early events that can support a prey to symbiont transition.

Conceptually, from the predator’s (host’s) point of view, the evolution of a mutualistic symbiosis requires them to benefit from a greater inheritable selective advantage by maintaining prey (symbionts) than by digesting them. Depending on the environment, energetic cost of maintaining the symbiont can be compensated by the by-products of photosynthesis such as exudated sugars or dioxygen molecule fuelling aerobic metabolism. We wished to test these scenarios using *T. thermophila* as a predator, *C. variabilis* and *S. elongatus* as photosynthetic eukaryotic and prokaryotic preys and *E. coli* as a non-photosynthetic prey.

In co-culture with *T. thermophila*, *E. coli* is rapidly phagocyted (Fig. S4) and can support growth of *T. thermophila* in a low-carbon media (Fig. 4E). Although fecal pellets containing fluorescent cells can be observed suggesting incomplete digestion (Fig. S4), this show that *E. coli* is fairly susceptible to phagocytosis and digestion by *T. thermophila* (Gurijala et Alexander, 1990; Matsui et al., 2000).

In low carbon media, *T. thermophila* derives only a slight growth advantage from the phagocytosis of *S. elongatus* in the dark that is enhanced in the light (Fig. 4C), showing a slight positive effect of photosynthesis that can be due to a better nutrient supply from cyanobacteria cells or by the production of exudates. *S. elongatus* is known to be digested by numerous groups of predatory unicellular eukaryotes (Christaki et al., 1999; Dillon et Parry, 2009; Dolan et Šimek, 1998), although it is a relatively poor-quality food source (Apple et al., 2011). The amount of exocytosed autofluorescent faecal pellets containin*g* undigested *S. elongatus* cells shows that it is partially resistant to digestion (Fig. 2C, D), explaining the relatively modest growth advantage provided by comparison to *E. coli*.

In contrast, *T. thermophila* appears unable to benefit from phagocytosis of *C. variabilis*, either by digestion or by the use of exudates (Fig. 4B and D). Indeed, a significant reduction in growth either in the light or in the dark indicates the presence of a cost (Fig. 4D). Species of Chlorella can be established as symbiont in different hosts, including ciliates. They are known to escape from the digestive vacuole into the cytoplasm to form a perialgal vacuole (or symbiosome) (Kodama et Sumita, 2022). Although the precise mechanism is unknown, *Chlorella* sp. could prevent fusion of the lysosome with the phagosome (Karakashian et Rudzinska, 1981) or acquire temporary resistance to lysosomal enzymes (Kodama et al., 2007).

The cost on *T. thermophila* growth by *C. variabilis* can have several causes. Chlorella is not known to release toxins, but a lethal effect of lectins produced by *Chlorella vulgaris* on mosquito *Aedes aegypti* larvae was observed (Cavalcanti et al., 2021). Intracellular accumulation without digestion of *C. variabilis* can inhibit *T. thermophila* growth or cell division. *C. variabilis* is progressively eliminated from *T. thermophila* cytoplasm as faecal pellets.

In sheer contrast to the modest or inexistent benefit provided by phagocyted phototrophs in low carbon media compared to *E. coli*, growth of *T. thermophila* was significantly stimulated in anoxia in the presence of either *C. variabilis* or *S. elongatus*. This effect was dependent on light, showing that photosynthesis was required. It was observed in conditions where most preys are intracellular (Fig. 5). Quantification of maximum oxygen production and consumption shows that the maximum number of phagocyted preys (40 *C. variabilis* and 160 *S. elongatus* per *T. thermophila* cell) are fully compatible with *T. thermophila* energy requirements in sub-atmospheric O_2_ concentration (Fig. 2, 3 and 7, Table 1).

This demonstrates that predation on photosynthetic cells can provide an immediate growth advantage caused by photosynthetically evolved oxygen in hypoxic environments. This advantage appears more consistent than the provision of sugars (or other carbon compounds) in carbon starved environments.

Evolution towards photosymbiosis can occur through different events such involving evasion from digestion by the prey (Lürling, 2021) or cell retention by the predator/host (Johnson, 2011). *Chlorella* species are known as photosymbionts of many different hosts (Hoshina et al., 2021; Matzke et al., 1990; Rajević et al., 2015; Zagata et al., 2016). *C. variabilis* cells are resistant to digestion by *T. thermophila,* but they slowly disappear from their cytoplasm. Evolution from prey to symbiont in this case can occur through cellular events repressing expulsion.

We further observed that the rate of prey phagocytosis decreases in hypoxia (Fig 6A and B) while that the cytoplasmic transit time increases (Fig. 6A, B). Similarly, In *T. pyriformis* hypoxia induces a generalized slowdown in metabolism, including the rate of phagocytosis (Skriver et Nilsson, 1978). Thus, by extending the prey retention time, hypoxia indirectly increases the benefit over time of oxygen intracellularly released by photosynthesis. Through genetic assimilation or accommodation (Braendle et Flatt, 2006; Kelly, 2019; Schneider et Meyer, 2017), these phenotypic changes in response to oxygen deficiency could promote genetic adaptation leading to photosynthetic endosymbiosis. Ecological observations further support that photosymbiosis occurs in hypoxia. In a stratified freshwater pond, 96% of ciliated protozoa living below the oxic-anoxic limit were found to host algae (Finlay et al., 1996). Oxygen produced by photosynthetic Symbiodiniaceae was also suggested to improve survival of *Anthopleura elegantissima* and *Anemonia viridis* hosts in hypoxic environments (Malcolm et Brown, 1977; Rands et al., 1992).

In aquatic environment, protozoa predators like ciliates are abundant and compete for feeding on large numbers of other microorganisms (Aijaz et Koudelka, 2017). The intracellular presence of phagocyted photosynthetic preys, even transiently, can provide an extended phenotype allowing expansion of the ecological niche into hypoxic hunting zones, and possible refuge from competitors and predators (Finlay et al., 1996; Gavelis et Gile, 2018; Quevarec et al., 2024).

Even in long evolved photosymbiotic organisms living in normoxic conditions, such as algae, mitochondria are able to use directly the photosynthetically evolved oxygen (Lavergne, 1989). Plants exposed to hypoxic stress by flooding also survive better in the light than in the dark suggesting a protective role of photosynthetic oxygen (León et al., 2021; Mommer et Visser, 2005).

Dioxygen can diffuse across membranes and is readily useable for respiration while photosynthetic sugars require dedicated transporters to cross symbiont and host membranes and require appropriate metabolic enzymes (Cenci et al., 2017). This makes O_2_ a more parsimonious driving force to stabilize early events leading to photosymbiosis, sugar supply probably resulting from further molecular and cellular exaptations (Gould et Vrba, 1982). Hypoxic environments are very common, can occur transiently, and were probably prevalent during a large part of life history. Indeed, while oxygenic photosynthesis evolved at least 3 Byr ago, it took around 1.6 Byr before the apparition of primary endosymbiosis at a time when oxygen was still scarce and its concentration highly variable, but where the terminal oxygen reductases that enable oxygen utilization already existed for a long time in prokaryotes (at least 3 Byr ago) and then eukaryotes via the mitochondria (2.2 Byr ago) (Degli Esposti et al., 2019; Falconet, 2012; Jabłońska et Tawfik, 2021). All in all, we propose that oxygen is a primary selective pressure for photosymbiosis evolution.

## Material and methods

### Organisms and population maintenance

The experiments were carried out with the *T. thermophila* strain CU 427 (Fig. 1A), obtained from Dr Mochizuki (CNRS – IGH, Montpellier). The ciliate *T. thermophila* is a strictly aerobic, mobile unicellular organism with an oral apparatus and a cytoproct for phagocytosis of prey and elimination of phagocytosis waste, respectively. As it is able to grow by osmotrophy, *T. thermophila* were routinely cultured in Neff medium (0.5% glucose, 0.25% peptone, 0.25% yeast extract and 33.3 µM FeCl_3_) (D. M. Cassidy-Hanley, 2012) at 30°C with shaking*. S. elongatus* UTEX 2973 (Fig. 1B) was routinely cultured in BG11 medium (Stanier et al., 1979) at 28°C in light (30 µEinstein.m^-2^.s^-1^) with shaking. *C. variabilis* NC64A (Fig. 1C) was cultured in TAP medium (Gorman et Levine, 1965) at 28°C in light (30 µEinstein.m^-2^.s^-1^) with shaking. TAP medium is supplemented with 700 µg. mL^−1^ ampicillin and 200 µg.mL^−1^ cefotaxime (Mustapa et al. 2016). *E. coli* strain MG1655 sod::gfp with the *sod* gene interrupted by the *gfp* gene was grown in LB media.

### Growth of Tetrahymena thermophila

We performed a standard curve to estimate the number of individuals as a function of the optical density (O.D.) measured with the cell density meter model 40 (Fisher Scientific) (λ = 600 nm) (Fig. S1). The number of individuals was measured using a Malassez counting chamber by microscopy.

Co-cultures of *T. thermophila* with preys *(S. elongatus*, *C. variabilis* or *E. coli)* was performed in rich Neff medium, in modified PP minimum medium (33.3 μM FeCl_3_, 0.25% proteose peptone) (D. M. Cassidy-Hanley, 2012) allowing very low growth or in minimum TAP medium (Gorman et Levine, 1965). The ‘Carbon’ experiments were carried out in Hungate tubes at 30°C with shaking in the dark or in the light (150 µEinstein.m^-2^.s^-1^) (Fig. 4). To test the advantage provided by oxygen from phagocytosed phototrophs, we co-cultured *T. thermophila* in Neff medium at 30°C under shaking with the different preys (*S. elongatus*, *C. variabilis* or *E. coli)* with oxygen or in anoxic sealed Hungate tubes in the dark or in the light (150 µEinstein.m^-2^.s^-1^) (Fig. 5B – D). Growth in low oxygen content was maintained by a massic gas mixer Pegas 4000 MF (Columbus Instruments) with a constant flux of 25 ml.min^−1^ using a mix of pure N_2_ and synthetic air (20% O_2_ and 80% N_2_). Concentration of O_2_ in the continuously flushed Hungate tubes was measured with a Mettler-Toledo M700 recorder equipped with an O_2_ module ppb 4700 and a nanomolar range sensitive Inpro 6900 O_2_ probe calibrated with two points 0 and saturated air (237µM O_2_ dissolved at 30°C). All cultures are in axenic conditions in Neff medium (Fig. 5A).

Microscopy and video-microscopy (Fig. 1 and Movie S1) were performed using an inverted Nikon Eclipse Ti microscope and a Motic B3 Professional microscope supplemented with a Moticam 4000 camera.

### Oxygen production and consumption by the different species

O_2_ consumption and production activities were measured polarographically with a Clark-type electrode in a stirred volume of 1.8 ml of corresponding medium saturated with air (237 µM O_2_ at 30 °C) (Fig. S5). Activities are expressed in nmol O_2_.min^-1^.mg^-1^ proteins (Fig. 7) (proteins content from whole cells were measured with Bradford solution + 0.1M NaOH using BSA as standard) or in 10^-^-^7^ nmol O_2_.min^-1^.cell^-1^ (Table 1) using standard curves (Fig. S1). Samples containing phototrophs were illuminated at light intensity of 100 µEinstein.m^-2^.s^-1^. All experiments were carried out at 30°C.

### Phagocytosis dynamics followed by flow cytometry

We measured the predation level of *T. thermophila* on the prey *S. elongatus*, *C. variabilis* or *E. coli* - GFP using flow cytometry with the measure of internal fluorescence accumulated in *T. thermophila* over time (Fig. 2, 3, 6, S2, S3 and S4). Thanks to the red autofluorescence of the phototrophs due to their photosynthetic pigments (Fig. 1F), we were able to follow timely and quantitatively the phagocytosis process knowing the level of fluorescence of the free partners. In addition, after exocytosis by cytoproct, we were also able to determine the number of preys in each dropping over time. For flow cytometry, the scatter density plots obtained (small-angle scattering FSC versus wide angle scattering SSC signals) were gated on the population of interest, filtered to remove multiple events and then analysed for the autofluorescence intensity of the phototrophs (filter 655/LP) or the GFP fluorescence (filter 525/30) of *E. coli* strain MG1655 sod::gfp. Samples were run in the low-pressure mode and a total number of 300-500K events were collected per sample. Data were acquired with a S3e cells sorter (Bio-rad) using 488 and 561 nm lasers and were analysed and plotted using FlowJo v10.6.

### Statistical analysis

Before the analysis, we square root-transformed data of O.D. or number of *T. thermophila*/µl for the growth curves in order to obtain normality and homoscedasticity of the data (Fig. 4 and 5). For *T. thermophila – E. coli* and *T. thermophila* – *S. elongatus* in modified PP minimum medium experiments we used raw data. We used a Generalized Additive Mixed Model (GAMM) with R software (R Core Team, 2013) and the Mgcv package (Wood et al., 2016) to analyze and compare the curves of several experiments with gaussian distribution. No overdispersion of the data was observed. We analyzed the number of *T. thermophila* as a function of time (in hours) (a continuous variable), culture conditions (i.e., medium, prey, presence of oxygen) as fixed effects. Replicate were added as random effects. The smoothing was performed on the variable time in function of culture condition. A contrast matrix is created for pairwise comparison of each culture condition with glht function of multcomp package (Hothorn et al., 2008) for ‘carbon’ experiments with at least 4 different treatments (Table S1C and S2C). We used Kruskal-Wallis rank sum test with R software and the stats package (R Core Team, 2013) to compare the growth of *T. thermophila* in anoxia with phototrophs (*S. elongatus, C. variabilis*) or *E. coli* in light and dark (Fig. 5). Then, a post hoc pairwise comparison (Dunn’s test) with Holm correction was realized between each treatment with rstatix package (Kassambara, 2019).

## Acknowledgments

We thank Dr. K. Mochizuki, T. Noto and J. Saksouk (IGH CNRS Montpellier, France), Aaron P. Turkewitz (Chicago University, U.S.A.) for the kind gift of *Tetrahymena thermophila* strains and precious advices. We thank the Cyanobacteria team at LCB-Marseille (Prof. A. Latifi, V. Risoul, A. Scholivet, S. Champ) for the kind gift of *Synechococcus elongatus* and the use of equipments, Dr. B. Ezraty and Prof. L. Aussel for *E. coli* strains, and Dr. Y. Li-Beisson and P. Auroy-Tarrago (BIAM) for *Chlorella variabilis*. We also thank Prof. D. Aragnol and S. Ridaoui for preliminary experiments, we thank the students V. Faroux and Y. Allio for their help with some experiments. Our IAM team is also acknowledged for support and Dr. L. Espinosa for microscopy advices. We thank P. Notareschi and A. Grossi for 3D printing accessories. This project PHOCEE N° 21-CE20-0035 to CR and GB was financially supported by the ANR agency and by IM2B (Institut de Microbiologie, Bioénergies et Biotechnologie, Aix-Marseille University) for a fellowship to RB.

## Author contributions

LQ, CR and GB conceptualized the study. LQ, RB, CR and GB designed the experiments. LQ, RB and GB performed the experiments and collected the data. LQ and GB performed the statistical analysis. LQ, CR and GB wrote the manuscript. All authors read, comment and approved the final manuscript.

