## Supplementary figures and tables for "Oxygen as a primary selective pressure for photo-endosymbiosis evolution"

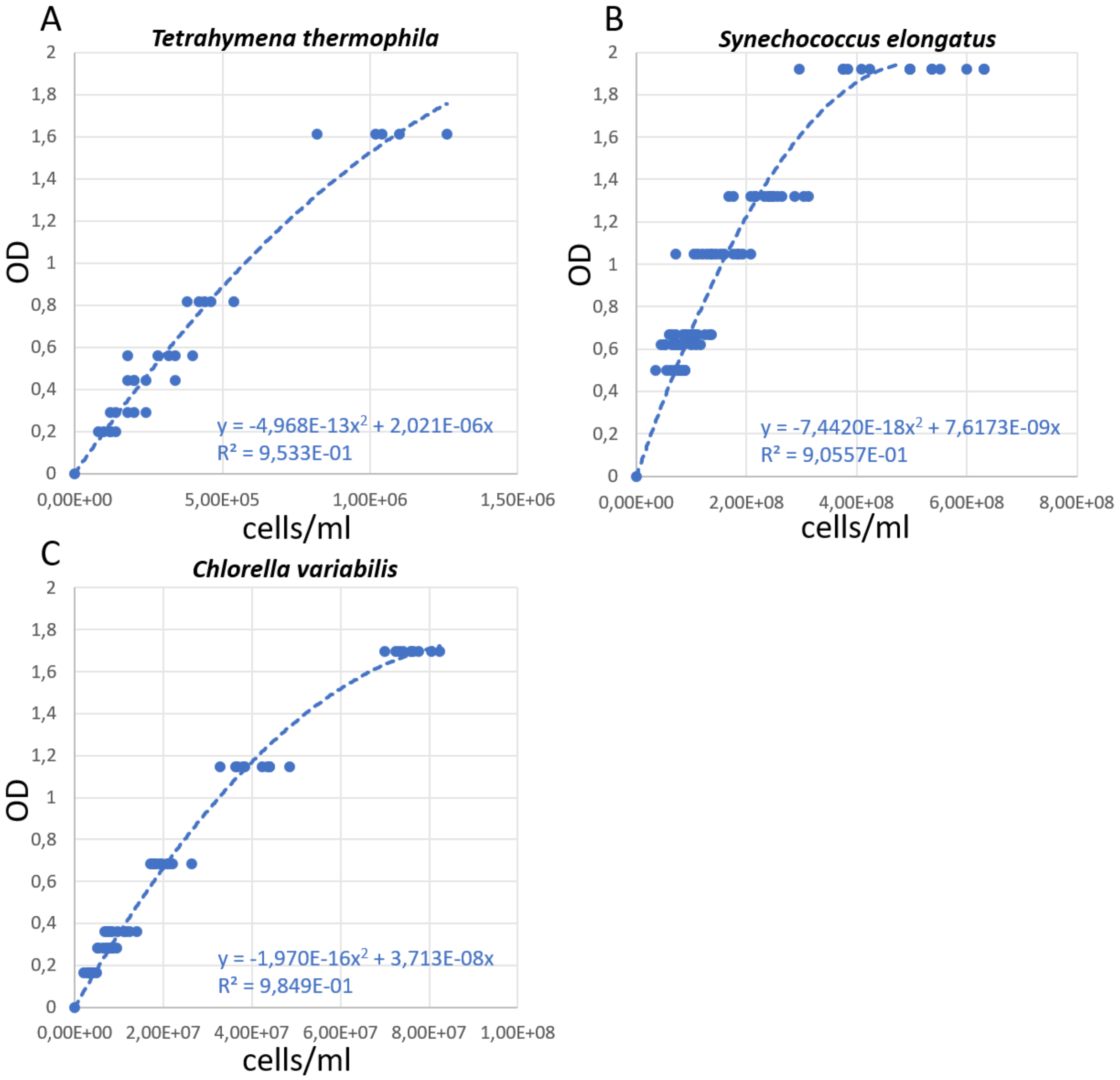


**Figure S1**: Standard curves for **A.** *T. thermophila*, **B.** *S. elongatus* and **C.** *C. variabilis* representing the optical density (O.D., $\lambda = 600$ nm ± 20 nm) as a function of the number of individuals counted on a Malassez counting chamber ($5 \leq n \leq 10$ for each concentration).

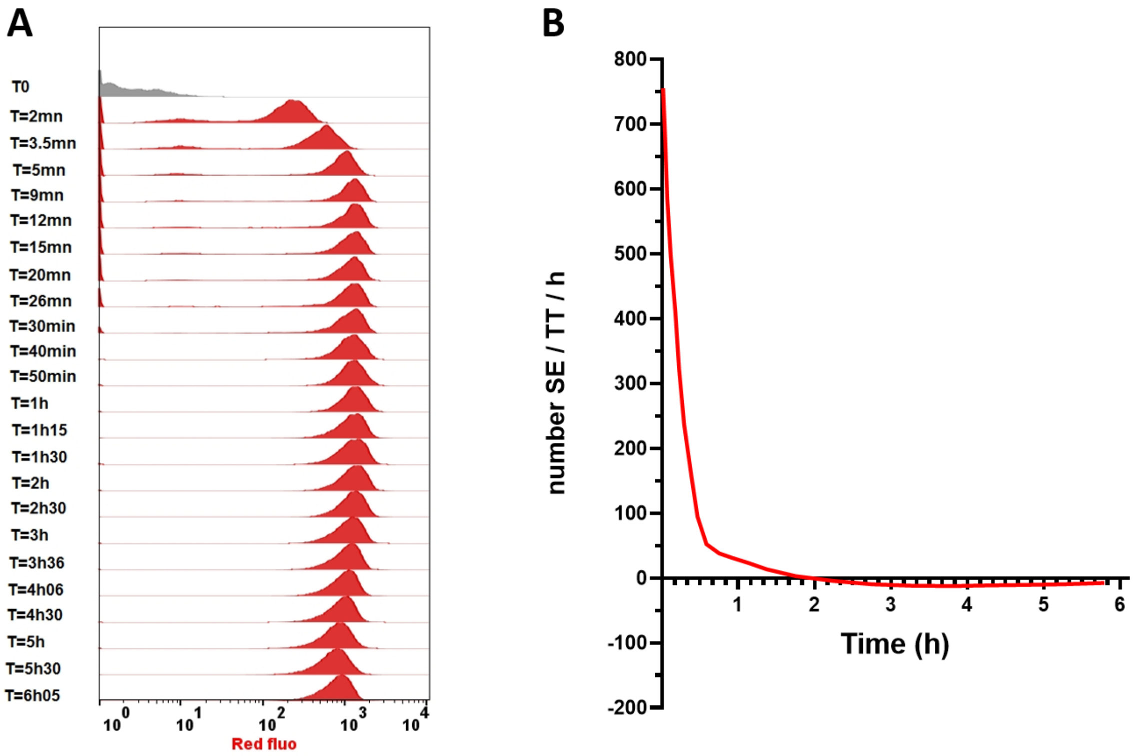


**Figure S2**: **Phagocytosis of cyanobacteria *S. elongatus* (SE) by the ciliate *T. thermophila* (TT). A)** Histograms of the intensity of red fluorescence of TT accumulated by phagocyting autofluorescent SE as a function of time measured by flow cytometry. X axis represents the intensity of TT red fluorescence in log scale. Y axis is the mode of number of events measured in the gated population of TT. **B)** First derivative kinetic of data in Fig. 2B representing the number of SE phagocyted/TT/hour as a function of time showing the fastest phagocytosis at the beginning and a decreasing kinetic over time until negative values corresponding to the digestion process. Maximum rate at the beginning reached about 750 SE phagocyted /TT/hour.

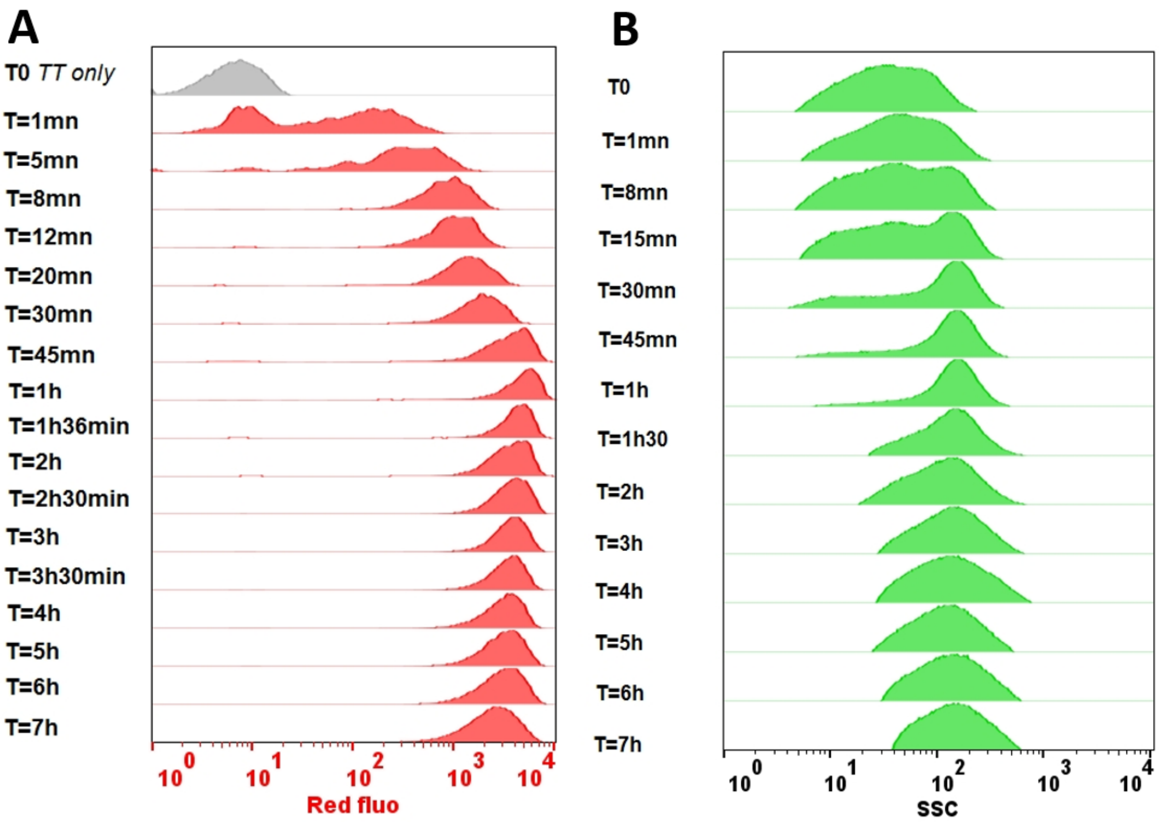


**Figure S3: Phagocytosis of *C. variabilis* (CV) by *T. Thermophila* (TT). A)** Histograms of the intensity of red fluorescence of TT accumulated by phagocyting autofluorescent CV as a function of time. X axis represents the intensity of TT red fluorescence in log scale. Y axis is the mode of number of events measured in the gated population of TT. **B)** Histograms of the intensity of side scatter SSC signal gated on the population of free CV showing the size selectivity of TT which first phagocytes medium and small size CV, larger CV remaining not phagocyted (see also Fig. 3A).

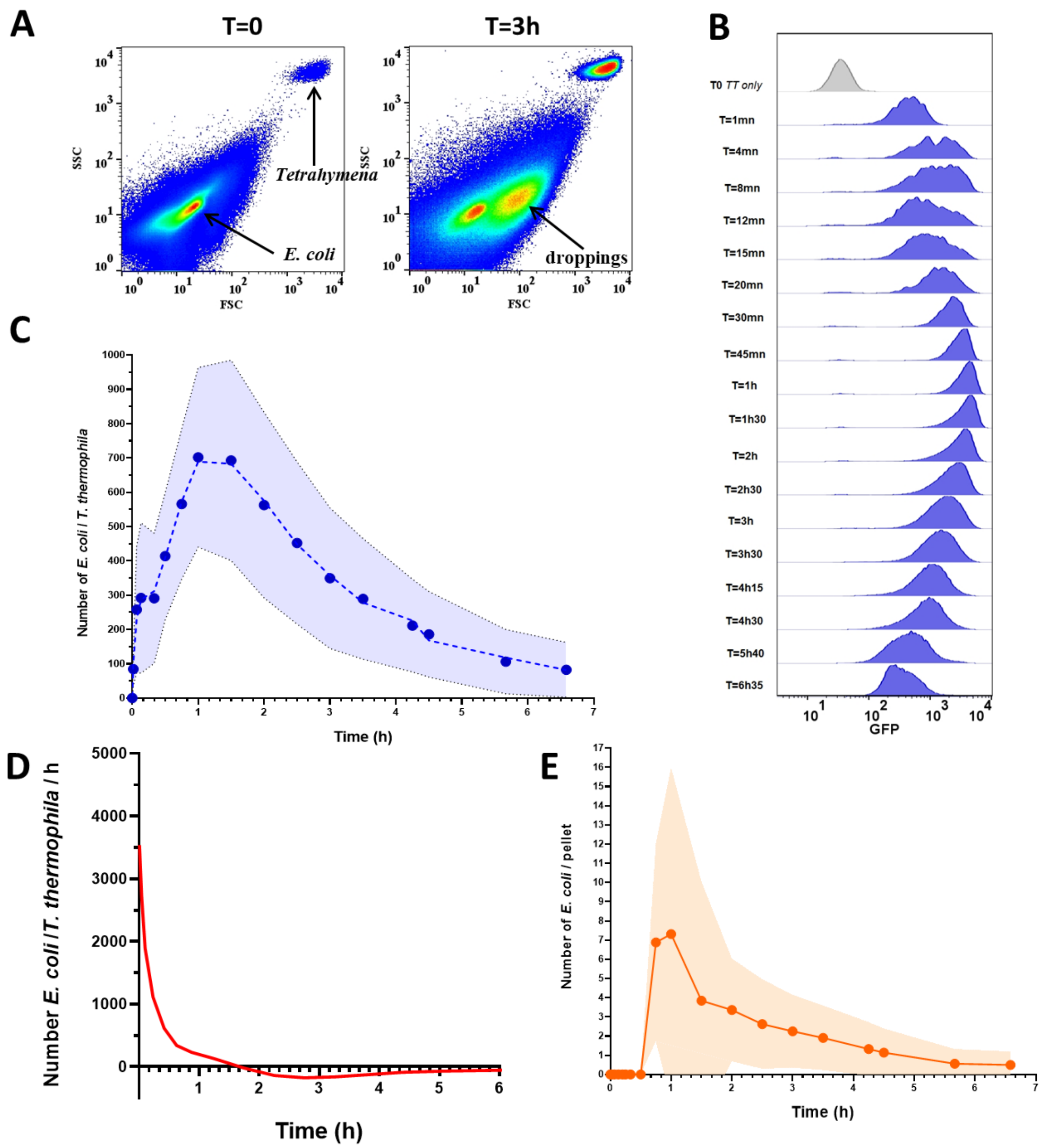


**Figure S4: Phagocytosis of *E. coli*-GFP (EC) by *T. thermophila* (TT) A)** Analysis of populations of EC-GFP (EC) mixed with TT in oxic conditions at 30°C (initial ratio of 10.000 EC/TT). Flow cytometry scatter density plots (SSC versus FSC) showing **TT**, the smaller diffraction of the EC cells and the apparition of a density region corresponding to the formation of TT droppings made of EC-GFP as a function of time. Color code from blue to red indicates low to high relative density of populations respectively, in a constant number of total events per time point (n = 3 x $10^6$). **B)** Histograms of the intensity of GFP fluorescence of TT accumulated by phagocyting EC-GFP as a function of time. X axis represents the intensity of TT green fluorescence in log scale and Y axis is the mode of number of events measured in the gated population of TT. **C)** Kinetics of the number of EC phagocyted per **TT** during fast phagocytosis (first phase) and digestion (second decreasing phase) using the green fluorescence of EC. Shaded area corresponds to the standard deviation of fluorescence mean. **D)** First derivative kinetic of data in **C)** representing the number of EC phagocyted/TT/hour as a function of time showing the fastest phagocytosis at the beginning and a decreasing kinetic over time until negative values corresponding to the digestion process. Maximum rate at the beginning reached about 3500 phagocyted EC/TT/hour. **E)** Kinetics of droppings formation with the number of EC-GFP per dropping as a function of time.

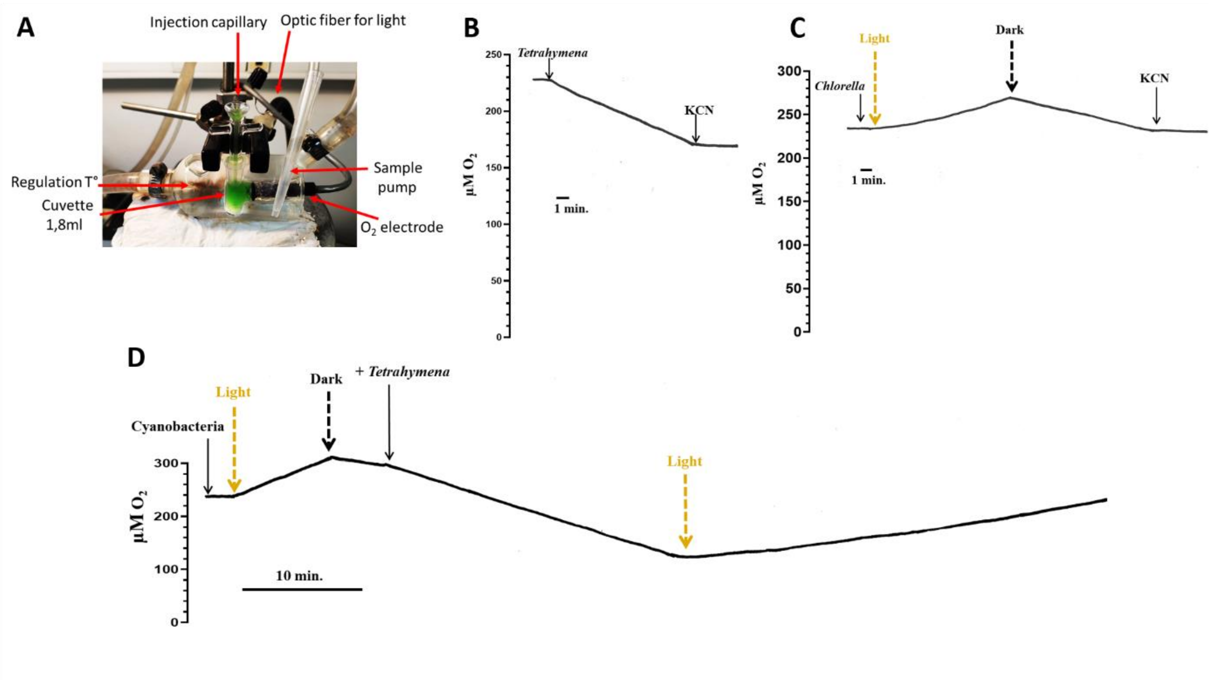


**Figure S5: $O_2$ consumption by *T. thermophila* (TT) and $O_2$ production by the phototrophs *S elongatus* (SE) and *C. variabilis* (CV)**. A) Device used to measure $O_2$ level in aqueous solution (TAP or BG11 buffer for **CV** or **SE**, respectively) using a Clark type electrode, a closed 1.8 ml temperature-controlled cuvette with an injection capillary and an optic fiber to illuminate the sample when needed with white lamp (saturating intensity between 50-100 µEinstein/m$^2$/s). **B)** $O_2$ consumption by **TT** cells (43 nmol $O_2$/min/mg prot. see Fig. 7) completely inhibited by KCN (1 mM). **C)** Production of $O_2$ by CV in the light and $O_2$ consumption in total darkness (see Fig. 7 for quantification). **D)** $O_2$ production by SE in the light and $O_2$ consumption in the total darkness (see Fig. 7 for quantification). Adding **TT** feeding on **SE** for 30 min does not impede $O_2$ production by **SE** inside **TT** when sample is illuminated for an additional 30 min, indicating that phagocyted cyanobacteria by TT are still able to produce $O_2$ at a similar rate.

**Table S1:** Effects of (a) culture conditions (minimum TAP medium or rich Neff medium with or without *S. elongatus* (SE)) and (b) time (EDF: effective degrees of freedom) on the growth of *T. thermophila* (TT). A pairwise comparison (c) was carried out. Ns: not significant; *, P < 0.05; **, P < 0.01; ***, P < 0.001.

**Parametric coefficients:**

| a) | Estimate | Std. Error | t value | Pr(>\|t\|) | |
|---|---|---|---|---|---|
| (Intercept) | 21.315 | 0.427 | 49.913 | $< 2 \times 10^{-16}$ | *** |
| Neff / TT+SE | -0.723 | 0.603 | -1.197 | 0.232 | ns |
| TAP / TT | -16.321 | 0.623 | -26.164 | $< 2 \times 10^{-16}$ | *** |
| TAP / TT+SE | -13.709 | 0.641 | -21.377 | $< 2 \times 10^{-16}$ | *** |

**Approximate significance of smooth terms:**

| b) | edf | Ref.df | F | p-value | |
|---|---|---|---|---|---|
| s(time): Neff / TT | 6.357 | 6.357 | 64.905 | $< 2 \times 10^{-16}$ | *** |
| s(time): Neff / TT+SE | 5.458 | 5.458 | 72.329 | $< 2 \times 10^{-16}$ | *** |
| s(time): TAP / TT | 1.000 | 1.000 | 0.348 | 0.555 | ns |
| s(time): TAP / TT+SE | 2.889 | 2.889 | 22.172 | $< 2 \times 10^{-16}$ | *** |

**Pair-wise comparison:**

| c) | Estimate | Std. Error | z value | Pr(>\|z\|) | |
|---|---|---|---|---|---|
| Neff / TT+SE - Neff / TT | -0.722 | 0.604 | -1.197 | 0.628 | ns |
| TAP / TT - Neff / TT | -16.321 | 0.624 | -26.164 | < 0.001 | *** |
| TAP / TT+SE - Neff / TT | -13.709 | 0.641 | -21.377 | < 0.001 | *** |
| TAP / TT - Neff / TT+SE | -15.599 | 0.624 | -25.015 | < 0.001 | *** |

| | | | | | |
|---|---|---|---|---|---|
| TAP / TT+SE - Neff / TT+SE | -12.986 | 0.641 | -20.258 | < 0.001 | *** |
| TAP / TT+SE - TAP / TT | 2.612 | 0.660 | 3.958 | < 0.001 | *** |

**Table S2:** Effects of (a) culture conditions (minimum TAP medium or rich Neff medium with or without *C. variabilis* (CV)) and (b) time (EDF: effective degrees of freedom) on the growth of *T. thermophila* (TT). A pairwise comparison (c) was carried out. Ns: not significant; *, $P < 0.05$; **, $P < 0.01$; ***, $P < 0.001$.

**Parametric coefficients:**

| a) | Estimate | Std. Error | t value | Pr(>\|t\|) | |
|---|---|---|---|---|---|
| (Intercept) | 21.467 | 1.327 | 16.183 | $< 2 \times 10^{-16}$ | *** |
| Neff / TT+CV | -0.954 | 1.865 | -0.512 | 0.609 | ns |
| TAP / TT | -15.865 | 1.829 | -8.676 | $< 2 \times 10^{-16}$ | *** |
| TAP / TT+CV | -15.161 | 1.836 | -8.259 | $4.11 \times 10^{-16}$ | *** |

**Approximate significance of smooth terms:**

| b) | edf | Ref.df | F | p-value | |
|---|---|---|---|---|---|
| s(time): Neff / TT | 6.241 | 6.241 | 270.405 | $< 2 \times 10^{-16}$ | *** |
| s(time): Neff / TT+CV | 5.994 | 5.994 | 245.674 | $< 2 \times 10^{-16}$ | *** |
| s(time): TAP / TT | 1.000 | 1.000 | 0.165 | 0.684 | ns |
| s(time): TAP / TT+CV | 3.208 | 3.208 | 31.705 | $< 2 \times 10^{-16}$ | *** |

**Pair-wise comparison:**

| c) | Estimate | Std. Error | z value | Pr(>\|z\|) | |
|---|---|---|---|---|---|
| Neff / TT+CV - Neff / TT | -0.954 | 1.865 | -0.512 | 0.956 | ns |
| TAP / TT - Neff / TT | -15.865 | 1.829 | -8.676 | $< 10^{-6}$ | *** |
| TAP / TT+CV - Neff / TT | -15.161 | 1.836 | -8.259 | $< 10^{-6}$ | *** |
| TAP / TT - Neff / TT+CV | -14.911 | 1.817 | -8.204 | $< 10^{-6}$ | *** |
| TAP / TT+CV - Neff / TT+CV | -14.207 | 1.825 | -7.786 | $< 10^{-6}$ | *** |
| TAP / TT+CV - TAP / TT | 0.704 | 1.787 | 0.394 | 0.979 | ns |

**Table S3:** Effects of (a) culture conditions (light or dark; with or without *S. elongatus* (SE)) and (b) time (EDF: effective degrees of freedom) on the growth of *T. thermophila* (TT) in modified minimum PP medium (PP). Ns: not significant; *, $P < 0.05$; **, $P < 0.01$; ***, $P < 0.001$.

**Parametric coefficients:**

| a) | Estimate | Std. Error | t value | Pr(>\|t\|) | |
|---|---|---|---|---|---|
| (Intercept) | 51.045 | 8.555 | 5.967 | 3.94e-09 | *** |
| PP+light / TT+SE | 49.182 | 12.096 | 4.066 | 5.36e-05 | *** |
| PP+dark / TT+SE | 7.128 | 12.096 | 0.589 | 0.556 | ns |

**Approximate significance of smooth terms:**

| b) | edf | Ref.df | F | p-value | |
|---|---|---|---|---|---|
| s(time): PP+light / TT | 5.153 | 5.153 | 13.14 | <2e-16 | *** |
| s(time): PP+light / TT+SE | 4.917 | 4.917 | 15.33 | <2e-16 | *** |
| s(time): PP+dark / TT+SE | 4.965 | 4.965 | 19.14 | <2e-16 | *** |

**Table S4:** Effects of (a) culture conditions (light or dark; with or without *C. variabilis* (CV)) and (b) time (EDF: effective degrees of freedom) on the growth of *T. thermophila* (TT) in modified minimum PP medium (PP). Ns: not significant; *, $P < 0.05$; **, $P < 0.01$; ***, $P < 0.001$.

**Parametric coefficients:**

| a) | Estimate | Std. Error | t value | Pr(>\|t\|) | |
|---|---|---|---|---|---|
| (Intercept) | 8.756 | 0.691 | 12.671 | < 2e-16 | *** |
| PP+light / TT+CV | -3.524 | 0.976 | -3.612 | 0.000324 | *** |
| PP+dark / TT+CV | -3.491 | 0.977 | -3.573 | 0.000376 | *** |

**Approximate significance of smooth terms:**

| b) | edf | Ref.df | F | p-value | |
|---|---|---|---|---|---|
| s(time): PP+light / TT | 3.896 | 3.896 | 29.146 | < 2e-16 | *** |
| s(time): PP+light / TT+CV | 2.351 | 2.351 | 2.538 | 0.149 | ns |
| s(time): PP+dark / TT+CV | 3.855 | 3.855 | 9.919 | 7.48e-07 | *** |

**Table S5:** Effects of (a) the presence of *E. coli* (EC) and (b) time (EDF: effective degrees of freedom) on the growth of *T. thermophila* (TT) in modified minimum PP medium (PP). ns: not significant; *, $P < 0.05$; **, $P < 0.01$; ***, $P < 0.001$.

**Parametric coefficients:**

| a) | Estimate | Std. Error | t value | Pr(>\|t\|) | |
|---|---|---|---|---|---|
| (Intercept) | 93.943 | 6.972 | 13.474 | < 2e-16 | *** |
| PP+light / TT+EC | 54.437 | 9.872 | 5.514 | 5.69e-08 | *** |

**Approximate significance of smooth terms:**

| b) | edf | Ref.df | F | p-value | |
|---|---|---|---|---|---|
| s(time): PP+light / TT | 3.825 | 3.825 | 19.81 | <2e-16 | *** |
| s(time): PP+light / TT+EC | 6.090 | 6.090 | 94.93 | <2e-16 | *** |

**Table S6:** Effects of (a) the oxygen concentration and (b) time (EDF: effective degrees of freedom) on the growth of *T. thermophila*. ns: not significant; *, $P < 0.05$; **, $P < 0.01$; ***, $P < 0.001$.

**Parametric coefficients:**

| a) | Estimate | Std. Error | t value | Pr(>\|t\|) | |
|---|---|---|---|---|---|
| (Intercept) | 0.0381 | 0.0672 | 0.567 | 0.572 | ns |
| 0.05% oxygen | 0.1926 | 0.0776 | 2.481 | 0.0138 | * |
| 0.1% oxygen | 0.1966 | 0.0776 | 2.534 | 0.0119 | * |
| 0.4% oxygen | 0.1987 | 0.0775 | 2.564 | 0.0110 | * |
| 1.4% oxygen | 0.3838 | 0.0823 | 4.663 | 5.22e-06 | *** |
| 5% oxygen | 0.5994 | 0.0824 | 7.271 | 5.27e-12 | *** |
| 21% oxygen | 0.7869 | 0.0774 | 10.170 | < 2e-16 | *** |

**Approximate significance of smooth terms:**

| b) | edf | Ref.df | F | p-value | |
|---|---|---|---|---|---|
| s(time): 0% oxygen (N2) | 1.828 | 1.828 | 1.201 | 0.365 | ns |
| s(time): 0.05% oxygen | 1.000 | 1.000 | 15.254 | 0.000123 | *** |
| s(time): 0.1% oxygen | 1.000 | 1.000 | 72.543 | < 2e-16 | *** |
| s(time): 0.4% oxygen | 3.106 | 3.106 | 138.045 | < 2e-16 | *** |
| s(time): 1.4% oxygen | 3.463 | 3.463 | 110.232 | < 2e-16 | *** |
| s(time): 5% oxygen | 3.699 | 3.699 | 295.663 | < 2e-16 | *** |
| s(time): 21% oxygen | 3.984 | 3.984 | 1131.154 | < 2e-16 | *** |

**Movie S1:** Time-lapse microscopy showing *T. thermophila* phagocytosing rapidly *C. variabilis* as shown by the initial high number of free red autofluorescent small *C. variabilis* cells and at the end (15 min) the low number of big red fluorescent *T. thermophila* cells.
